# SH3-domain mutations selectively disrupt Csk homodimerization or PTPN22 binding

**DOI:** 10.1101/2022.01.13.475555

**Authors:** Ben F. Brian, Frances V. Sjaastad, Tanya S. Freedman

## Abstract

The kinase Csk is the primary negative regulator of the Src-family kinases (SFKs, e.g., Lck, Fyn, Lyn, Hck, Fgr, Blk, Yes), phosphorylating a tyrosine on the SFK C-terminal tail that mediates autoinhibition. Csk also binds phosphatases, including PTPN12 (PTP-PEST) and immune-cell PTPN22 (LYP/Pep), which dephosphorylate the SFK activation loop to promote autoinhibition. Csk-binding proteins (e.g., CBP/PAG1) oligomerize within membrane microdomains, and high local concentration promotes Csk function. Purified Csk homodimerizes in solution through an interface that overlaps the phosphatase binding footprint. Here we demonstrate that Csk can homodimerize in Jurkat T cells, in competition with PTPN22 binding. We designed SH3-domain mutations in Csk that selectively impair homodimerization (H21I) or PTPN22 binding (K43D) and verified their kinase activity in solution. Disruption of either interaction, however, decreased the negative-regulatory function of Csk in cells. Csk W47A, a substitution previously reported to block PTPN22 binding, also impairs homodimerization. Csk H21I and K43D will be useful tools for dissecting the protein-specific drivers of autoimmunity mediated by the human polymorphism PTPN22 R620W, which impairs interaction with Csk and with the E3 ubiquitin ligase TRAF3. Future investigations of Csk homodimer activity and phosphatase interactions may reveal new facets of SFK regulation in hematopoietic and non-hematopoietic cells.

## Introduction

The protein tyrosine kinase Csk negatively regulates Src-family kinases (SFKs) by phosphorylating the C-terminal inhibitory tail to promote assembly of an autoinhibited state^1–3^. Unlike the SFKs, Csk lacks membrane-anchoring lipidation sites, and its activity is not regulated by phosphorylation of its kinase-domain activation loop^1^. Instead, Csk is recruited to membrane-anchored SFKs and activated allosterically via its SH2 domain, which binds SFK-phosphorylated motifs in CBP/PAG1 (Csk binding protein/phosphoprotein associated with glyco-sphingolipid-enriched microdomains), paxillin, Fak, the Dok family, and other tyrosine-phosphorylated proteins^1,4–7^. Csk has low intrinsic activity toward peptide substrates and requires a stable interaction through an extensive kinase-domain/kinase-domain binding interface for optimal SFK substrate phosphorylation^8^. Colocalization with the SFKs and high local concentrations are therefore key prerequisites for Csk-mediated suppression of cell signaling.

Like the SH2 domain, the SH3 domain of Csk is required for kinase activity and for mediating protein-protein interactions^9–11^. Primary ligands are PXXP-containing tyrosine phosphatases, including broadly expressed PTPN^12^ (PTP-PEST)12 and hematopoietic-cell PTPN22 (LYP in humans, Pep in mice)^10^. PTPN12 binds the SH3 domain of Csk via a canonical polyproline (PXXP) interaction (K_d_ ~8 μM). The SH3-binding peptide of PTPN22 has a more extensive footprint, wrapping around the SH3 domain to bind with 10-fold higher affinity (K_d_ ~0.8 μM)^12,13^. Along with Csk, PTPN12 and PTPN22 are critical modulators of homeostatic signaling and negative feedback. In addition to restraining SFK signaling directly by dephosphorylating the SFK activation loop, they reverse the phosphorylation of SFK substrates^10,12,14^.

In T lymphocytes the dominant SFK, Lck, phosphorylates and activates the downstream kinase Zap70 and ITAMs (immunoreceptor tyrosine-based activation motifs) within the T-cell antigen receptor (TCR) complex^15–17^. The activity of Lck is tuned to provide homeostatic function and inducible signaling upon TCR engagement. Csk and PTPN22 are both required to maintain this balance. PTPN22 dephosphorylates the activation loop of Lck, the ITAMs of the CD3ζ TCR subunit, and the activation loop of Zap70^18–20^, adding a second level of TCR signal inhibition to the direct activity of Csk on Lck. Dysregulated Lck/TCR/Zap70 signaling, either too strong or too weak, can lead to autoimmune disease^21^, and Csk or PTPN22 dysregulation can promote autoimmunity and cancer^22–25^. Impairment of PTPN22 binding to the SH3 domain of Csk by an arginine-to-tryptophan substitution at position 620 (R620W)^26,27^, is linked to autoimmune disease in humans and mice^28–31^.

The role of the wild-type (WT) PTPN22/Csk complex in T-cell and other immune-cell signaling is not yet clear^32^. In some reports, binding to Csk appears to sequester and functionally suppress WT PTPN22, and the R620W variant is a gain-of-function allele^32–36^. Others have found the opposite, that the complex with Csk promotes PTPN22 function by colocalizing it with substrates^10,37–39^. In this latter model, PTPN22 R620W is a loss-of-function allele. The contributions of cell-type and species differences to these disparate reports is unclear. Further complicating these analyses, R620W also blocks PTPN22 binding to the E3 ubiquitin ligase TRAF3. In addition to promoting NF-κB signaling^40,41^, TRAF3 normally has an adaptor-like function in binding and inhibiting Csk and PTPN22, possibly as a ternary complex^42^.

Csk homodimer formation could limit the abundance of WT PTPN22/Csk complexes and contribute to PTPN22 R620W autoimmune disease. Purified Csk protein homodimerizes in solution with low-to-moderate affinity (K_d_ ~10-20 μM^43^). The homodimer interface, mediated by symmetrical SH3-SH3 interactions that bridge two Csk molecules^44,45^, overlaps completely with the PXXP docking site for PTPN12^12,44^ and overlaps substantially (but not completely) with the extended binding site for PTPN22^13,43^. In cells, phosphorylated CBP/PAG1 can oligomerize, binding multiple molecules of Csk in supercomplexes localized to membrane microdomains or rafts^45^. At these high local concentrations, Csk homodimer interactions could compete with phosphatases for access to SH3-domain binding sites. However, Csk homodimerization and competition with PTPN22 in cells has not, to our knowledge, been investigated.

Here we report that Csk self-association in Jurkat T cells can compete with PTPN22 for binding to the SH3 domain of Csk. Using structural analysis, we identified a histidine-to-isoleucine substitution (H21I) that blocked co-immunoprecipitation of Myc- and HA-tagged Csk and increased co-immunoprecipitation of endogenous PTPN22. A lysine-to-aspartate substitution (K43D) in the SH3 domain of Csk blocked PTPN22 binding without disrupting Csk self-association. Neither mutation impaired the activity of purified Csk in solution, but both mutations decreased the suppressive function of Csk in TCR signaling. The tryptophan-to-alanine substitution (W47A), used previously to block PXXP-Csk interactions46-50, disrupted both PTPN22 binding and Csk homodimer formation. Together, our studies demonstrate that Csk can homodimerize in cells and restrict the formation of Csk/PTPN22 complexes. Our findings also suggest that asynchronous homodimerization and PTPN22 binding are both required for the full function of Csk in suppressing TCR signaling.

## Results

### Multiple molecules of epitope-tagged Csk co-immunoprecipitate from Jurkat-cell lysates

We investigated Csk self-association in Jurkat T cells by co-transfecting HA-tagged Csk (Csk^HA^) and Myc-tagged Csk (Csk^Myc^) and subjecting cell lysates to anti-Myc immunoprecipitation. Csk^HA^ was detected in Csk^Myc^ immuno-precipitates, suggesting interaction of Csk^HA^ and Csk^Myc^ proteins **(Fig. 1a)**. Blots of whole-cell and immuno-depleted lysates demonstrated the specificity of the HA and Myc antibodies and the efficiency of Myc immuno-precipitation **(Fig. 1b)**. Lack of detectable Csk^HA^ depletion from Myc-immunodepleted lysates (i.e., substoichiometric binding of Csk^HA^ to Csk^Myc^) is consistent with the moderately low affinity and fast on/off kinetics of dimerization in solution^43^. Other factors that could limit Csk^HA^ co-immunoprecipitation include competition of homodimerization with binding of endogenous PTPN22 and signal interference from other dimer pairings, such as Csk^Myc^/Csk^Myc^, Csk^HA^/Csk^HA^, and Csk^Myc^/endogenous Csk.

**Fig. 1.**
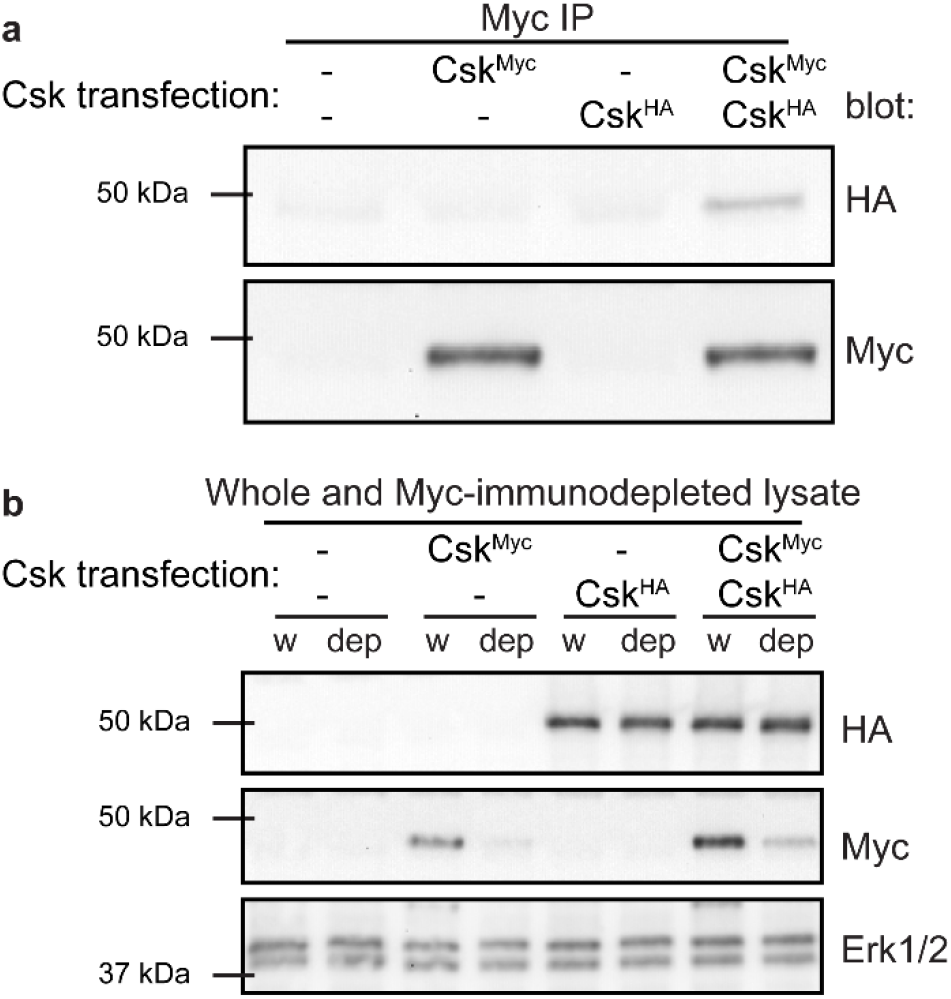
Csk self-associates when overexpressed in Jurkat cells. **(a)** Immunoblots of anti-Myc immunoprecipitates (IP) from transiently transfected Jurkat T cells. Transfection conditions are indicated, including epitope-tagged Csk constructs (Csk^Myc^, Csk^HA^) and empty pEF6 vector (-).Blots are representative of at least three independent experiments. **(b)** Immunoblots showing transfected Csk^Myc^ and Csk^HA^ and endogenous Erk1/2 loading control in whole-cell lysate (w) and Myc-immunodepleted lysate (dep) from transfected Jurkat cells; samples correspond to the immunoprecipitations above. Blots are representative of at least three independent experiments.

### An amino-acid substitution in the Csk SH3 domain impairs homodimerization and enhances PTPN22 binding

The Csk- and PTPN22-binding surfaces lie within the SH3 domain of Csk^13,43,51^ **(Fig. 2a)**. Although the footprints of these interfaces overlap substantially, we identified unique interactions via structural alignment of the dimer and extended PTPN22 binding surfaces from published crystal and NMR structures^13,44,51^. A single residue, H21, was situated in the symmetrical homodimer interface and outside the PTPN22 binding footprint **(Fig. 2b)**. Analysis with Pymol software suggested that most amino acids could adopt non-clashing rotamers if swapped into this position; β-branched isoleucine was an exception. We therefore generated H21I variants of Csk^Myc^ and Csk^HA^ and tested their effect on Csk co-immunoprecipitation.

**Fig 2.**
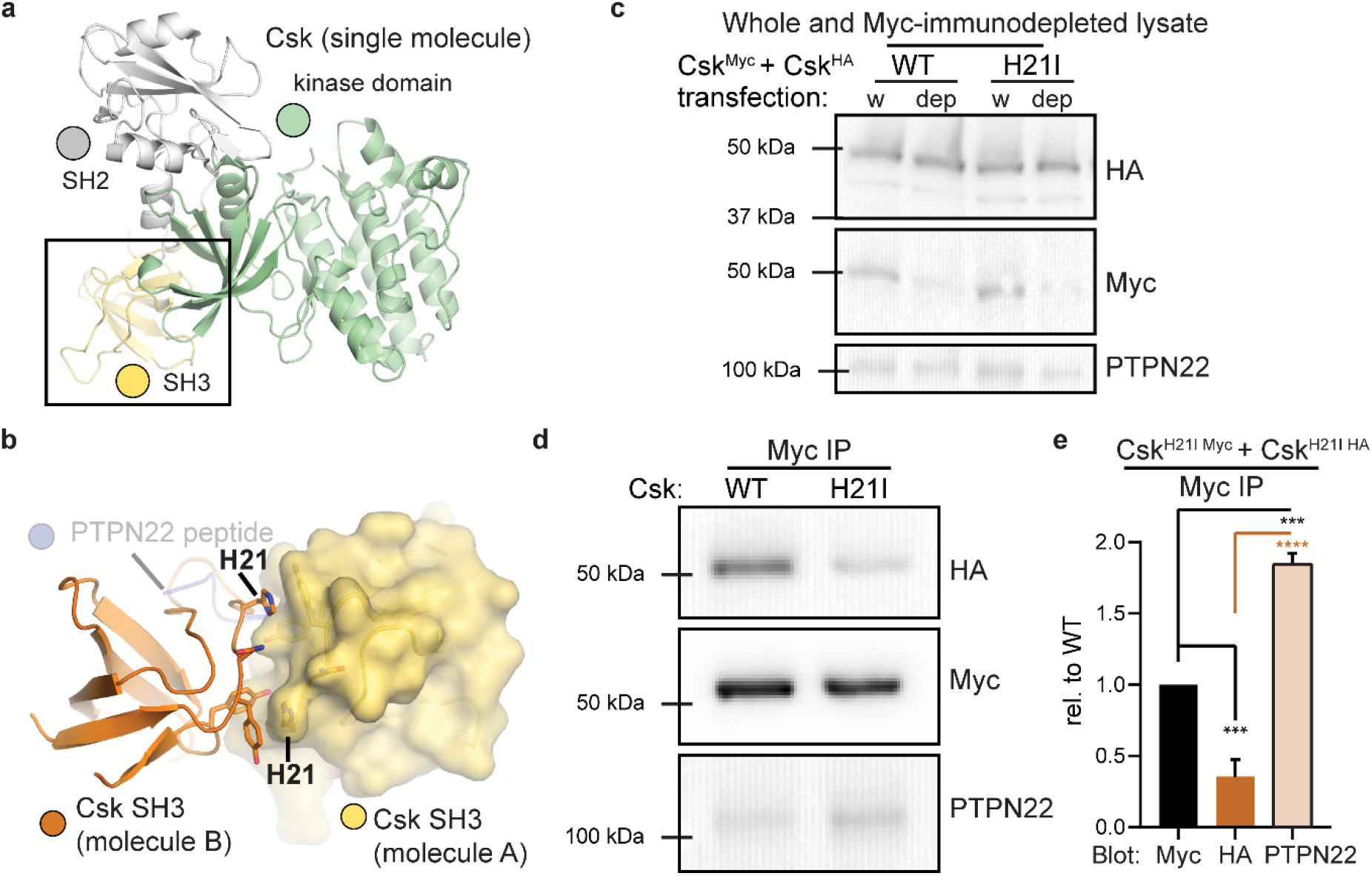
H21I substitution in the SH3 domain of Csk selectively disrupt self-association. **(a)** Position of the SH3 domain in full-length Csk (protein databank identifier (PDB) 1K9A^52^). **(b)** Csk residue H21 in the SH3/SH3 homodimer interface (PDB 1CSK^51^). The position of PTPN22 peptide binding to the SH3 domain was derived in Pymol using PDBs 1K9A^53^ 1CSK^51^, 1JEG^13^, 1QWE^44^, and 2P6X^54^. **(c)** Representative immunoblots of whole-cell or Myc-immunodepleted lysates from Jurkat cells transfected with Csk^HA^ and Csk^Myc^ constructs (both WT or both H21I). **(d)** Corresponding immunoblots of transfected Csk and endogenous PTPN22 in Myc immunoprecipitates. **(e)** Quantifications from immunoprecipitate blots, corrected for Csk^Myc^ pulldown in the same sample, shown relative to WT. Error bars: standard error of the mean (SEM), n=4 independent experiments. Significance: one-way ANOVA with Tukey’s multiple comparison test (Sig.ANOVA) ****p<0.0001, ***p=0.0001 or 0.0008.

We transiently transfected Jurkat cells with pairs of Csk^Myc^ and Csk^HA^ constructs, both either WT or H21I. Protein expression was not altered by H21I substitution, and the Myc-tagged construct was effectively immunodepleted **(Fig. 2c)**. H21I substitution disrupted Csk^HA^ co-immunoprecipitation with Csk^Myc^ (60 ± 20% less HA-tagged H21I than WT) **(Fig. 2d-e)**. This deficit was accompanied by increased co-immunoprecipitation of endogenous PTPN22 (80 ± 20% more in H21I-transfected cells than in WT-transfected cells). Together, these data suggest that Csk dimerization can compete with PTPN22 binding. This competition also reveals that the uniquely extended SH3-binding motif in the PTPN22 peptide^10^ cannot bind stably to the SH3 domain of Csk in the absence of canonical PXXP-motif docking.

Although a previous study probed the SH3 homodimer interface using point mutations^43^, the authors did not test the effect of H21I substitution on dimerization. We purified recombinant Csk WT and H21I from bacteria and assessed their mobility on a size exclusion column. As expected, purified Csk WT and H21I were each visible as a single SDS PAGE band of ~50 kDa, consistent with the 50.6 kDa molecular weight of full-length Csk **(Fig. 3a inset)**. Csk H21I, however, was retained longer than WT on a size exclusion column **(Fig 3a)**, reflecting a decrease in apparent molecular weight^44,53,54^. The Csk H21I retention volume was consistent with a monomeric species, confirming disruption of the homodimer interface **(Fig 3b)**^43,55,56^. The more rapid migration of WT Csk through the column corresponded to a molecular weight of ~70 kDa, smaller than a stable dimeric species of 101 kDa. This lower apparent molecular weight is consistent with a rapidly exchanging population of monomers and dimers after dilution of the sample in the column^43^. Together the co-immunoprecipitation of Csk^HA^ with Csk^Myc^ and the observation that a structure-guided point mutation selectively disrupts this interaction, in cells and in purified protein, lead us to conclude that Csk homodimerizes in Jurkat T cells.

**Fig. 3.**
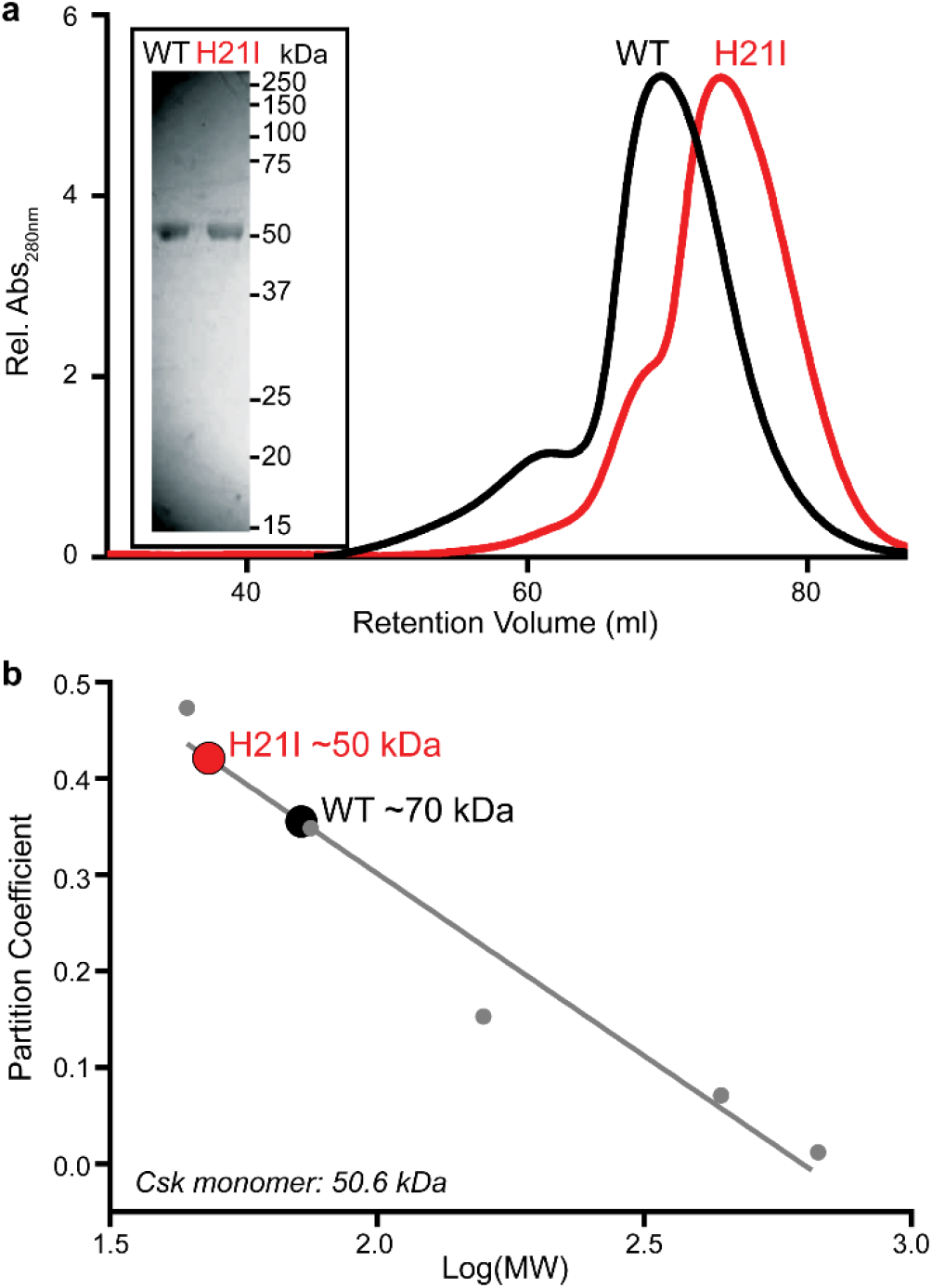
Mutation of the homodimer interface decreases the apparent molecular weight of Csk in solution. **(a)** Colloidal blue staining (reducing SDS PAGE, inset) and Superdex 200 size-exclusion column migration of purified, recombinant Csk WT and H21I. **(b)** The apparent molecular weight of each Csk variant in solution was approximated based on column partitioning of proteins with known molecular weights.

### An amino-acid substitution in the Csk SH3 domain selectively impairs PTPN22 binding

The homodimer interface in the Csk SH3 domain overlaps completely with the canonical PXXP-binding site shared by PTPN12 and PTPN22. A second point of interaction with Csk, however, is unique to PTPN22 (LYP in humans, Pep in mice)^43^. Structural modeling of this extended interaction suggested that K43 in the Csk SH3 domain uniquely participates in PTPN22 binding but not homodimer formation **(Fig. 4a)**. Since this positively charged residue forms a salt bridge with a nearby aspartate in PTPN22, we flipped the charge and minimized the degrees of rotameric freedom with a K43D substitution.

**Fig. 4.**
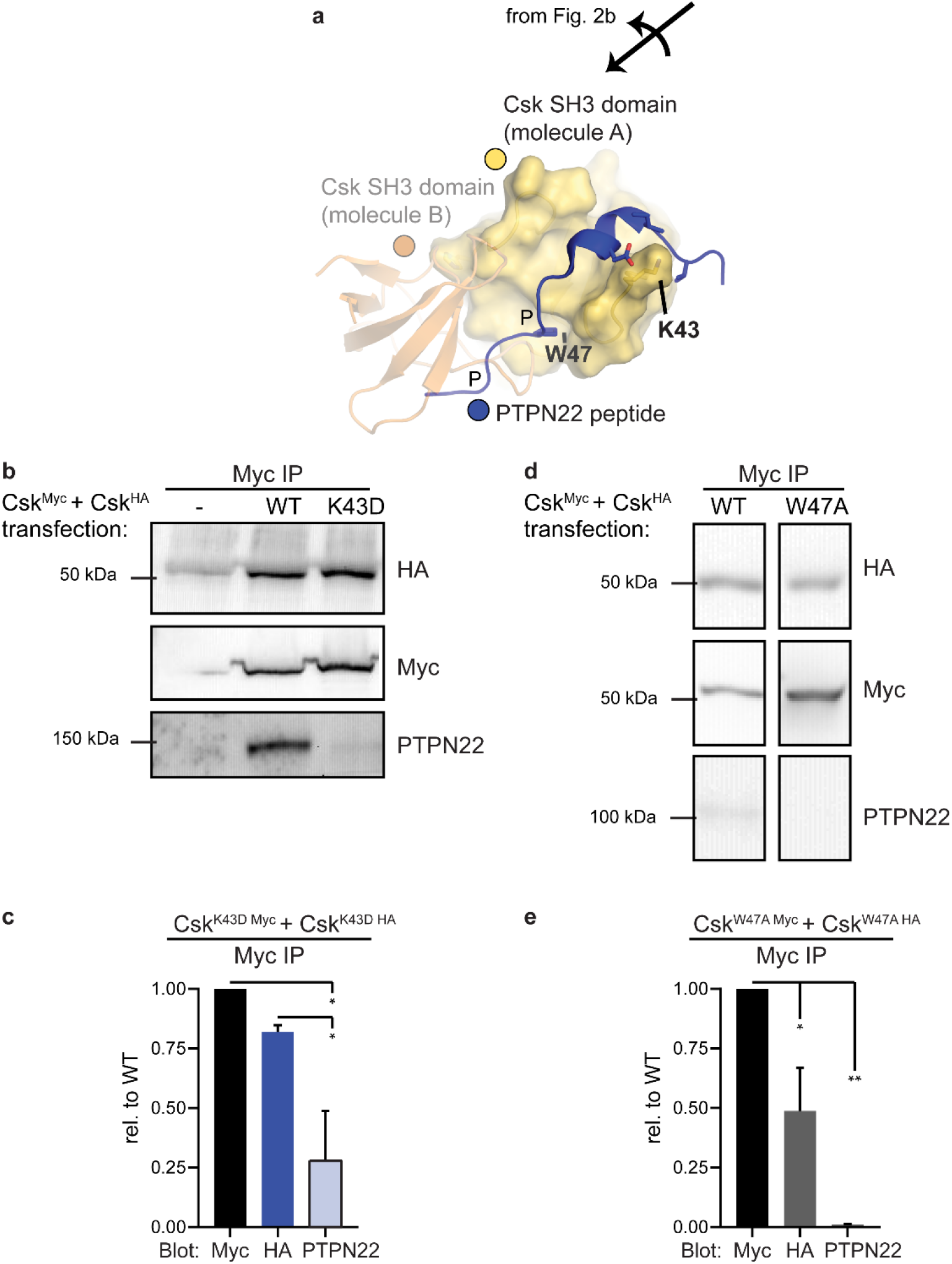
K43D substitution in the SH3 domain of Csk selectively disrupts PTPN22 binding but not self-association, whereas W47A substitution impairs both interactions. **(a)** Csk residue K43D in the secondary binding interface between PTPN22 and the SH3 domain of Csk. The location of the proline (P) residues within the peptide PXXP motif is shown, overlapping the dimer footprint; only one of the proline residues is engaged in this structure. Csk residue W47 is an essential component of the indicated proline binding pocket. **(b)** Representative immunoprecipitate blots from Jurkat cells transfected with Csk^HA^ and Csk^Myc^ constructs (both WT or both K43D). **(c)** Quantifications from immunoprecipitate blots, corrected for Csk^Myc^ pulldown in the same sample, shown relative to WT. Error bars: SEM, n=3. Sig.ANOVA *p=0.0133 or 0.0451. **(d)** Representative immunoblots of immunoprecipitates from Jurkat cells transfected with Csk^HA^ and Csk^Myc^ constructs (both WT or both W47A). Boxed images were cropped from non-adjacent lanes of the same blot; brightness/contrast corrections were applied to the whole blot prior to cropping. **(e)** Quantifications from immunoprecipitation blots, corrected as above. Error bars: SEM, n=4 independent experiments. Sig.ANOVA **p=0.0010, *p=0.0282.

Endogenous PTPN22 co-immunoprecipitated poorly with Csk K43D (70% ± 30% less than WT), without a significant secondary defect in Csk^HA^ co-immunoprecipitation **(Fig. 4b-c)**. The selectivity of the H21I and K43D substitutions for impairing Csk or PTPN22 co-immunoprecipitation, respectively, also demonstrate the adequate stringency of the co-immunoprecipitation protocol for detecting specific protein-protein interactions.

We also generated Myc- and HA-tagged constructs of Csk W47A, which has been used previously to ablate the binding of PXXP-containing ligands, including PTPN22, to the SH3 domain of Csk^46–50^. Residue W47 forms one of the proline binding pockets for the canonical PXXP interaction and lies within the homodimerization footprint **(Fig. 4a)**. Co-immunoprecipitation experiments confirmed that the Csk W47A substitution impaired both Csk/Csk and Csk/PTPN22 interactions **(Fig. 4d-e)**. This suggests that a defect in Csk homodimer formation could be a complicating factor in previous studies of PTPN22 and other SH3-binding proteins; going forward, the K43D mutation could be a more specific tool for disrupting Csk binding to PTPN22.

### H21I and K43D substitutions do not impair the kinase activity of Csk in solution but may alter cellular function

Full-length Csk WT, H21I, K43D, and K222R (a kinase-impaired lysine-to-arginine mutant)^3^ proteins were purified and tested for activity against an optimal substrate peptide^57^. Activity was measured using a continuous spectrophotometric assay coupling ATP hydrolysis to NADH oxidation and decreasing absorbance at 340 nm **(Fig. 5a)**^53^. Neither the H21I substitution nor K43D impaired kinase activity in solution **(Fig. 5b)**. As expected, Csk K222R was nearly kinase dead (94% ± 5% loss of activity compared to WT).

**Fig. 5.**
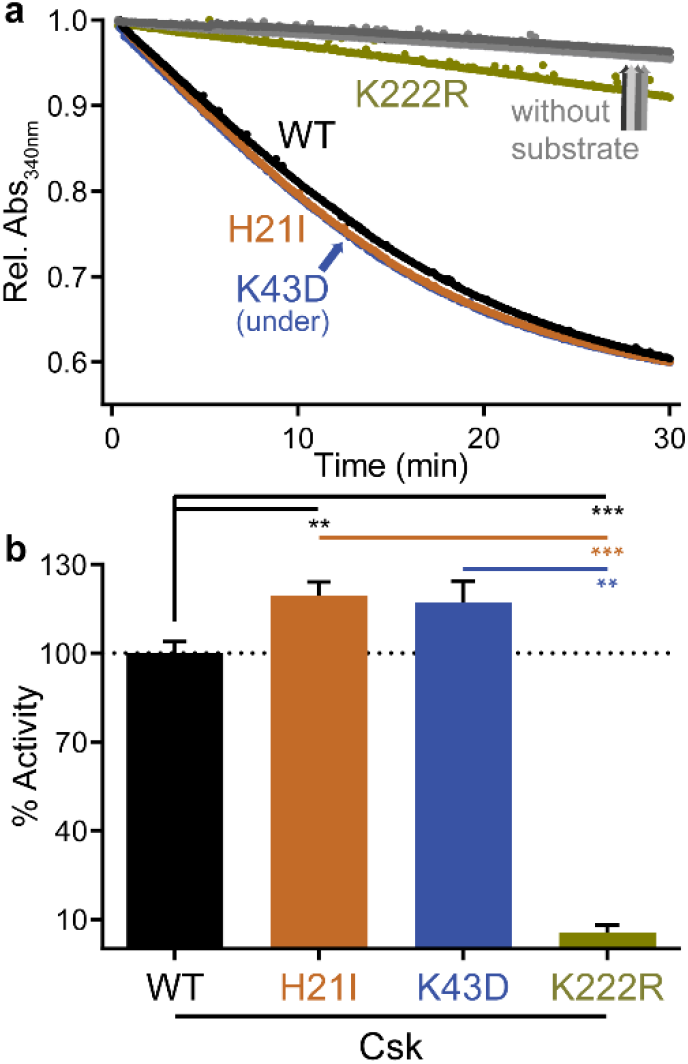
H21I and K43D SH3-domain substitutions do not impair the catalytic activity of Csk in solution. **(a)**Continuous spectrophotometric assay of peptide phosphorylation, in which ATP hydrolysis is coupled via pyruvate kinase and lactate dehydrogenase to NADH oxidation and loss of absorbance at 340 nm. Reactions were initiated by adding Csk (WT, H21I, K43D, or K222R) to the reaction mix. Control reactions without substrate peptide are also shown. **(b)** Linear velocities of the initial kinase reaction relative to WT. Error bars: SEM, n=4 independent experiments.

As an initial test of homodimer function in cell signaling, we measured the ability of Csk H21I to suppress Lck-dependent TCR triggering^16^. Signaling was initiated by treating Csk^Myc^ transient transfectants with the C305 antibody, which ligates the Jurkat-cell TCR^58^. After 2 min, we quenched signaling and performed intracellular staining for Myc (a proxy for Csk-construct expression) and phosphorylated Erk1/2 threonine 202/tyrosine 204 (pErk, a proxy for TCR-pathway activation^16^). We then performed flow cytometry to probe the effect of Csk expression on TCR signaling. In the heterogenous pool of transient transfectants, we observed a complete loss of pErk positivity in cells with the highest expression of WT Csk. **(Fig. 6a)**. As induction of pErk downstream of the TCR in a given cell is an all-or-none response^59^, quantification of the fractional content of pErk-positive (^+^) cells at each Csk dose was well described by a sigmoidal function. We therefore fit each data set to a dose-response curve to generate an apparent IC_50_ value, a relative measure of the Csk dose at which the frequency of pErk^+^ cells was reduced by 50% **(Fig. 6b)**. Of the constructs tested, WT Csk suppressed TCR signaling most efficiently **(Fig. 6c)**. At the other extreme, Csk K222R was unable to fully suppress TCR signaling, even at the highest expression levels. Csk H21I and Csk K43D both had more subtle but still significant functional defects, with apparent IC_50_ values increased relative to WT (20% ± 10% and 30% ± 20% higher, respectively) **(Fig. 6d)**. Together, these data suggest that blocking homodimerization or PTPN22 binding decreases the immunosuppressive function of Csk in cells, an effect that cannot be attributed to a simple loss of intrinsic catalytic function.

**Fig. 6.**
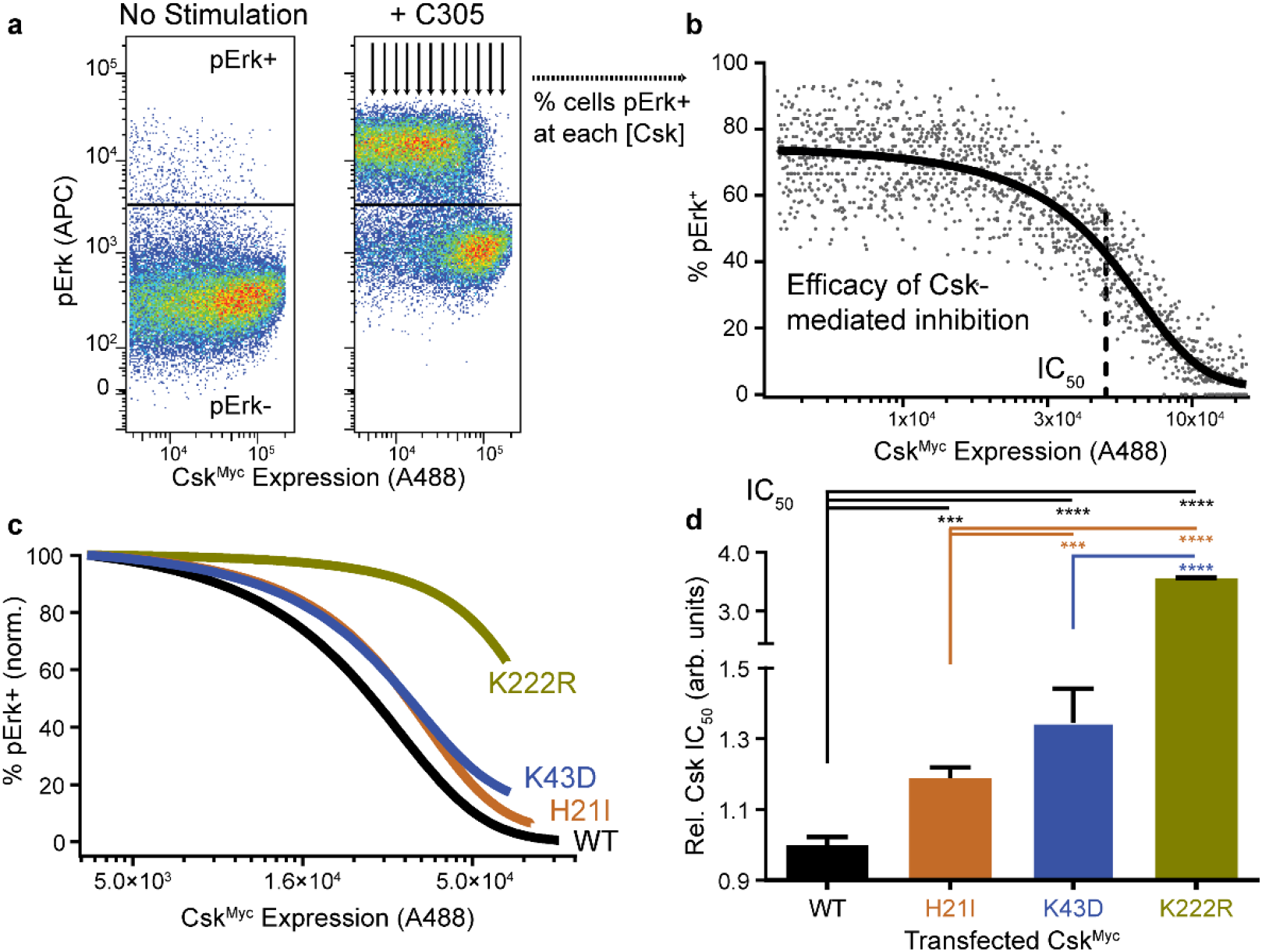
Loss of Csk dimerization may impair its function in T cells. **(a)** Jurkat T cells transiently transfected with Csk^Myc^ and rested (left) or TCR-stimulated for 2 min using C305 antibody and stained for intracellular Myc and pErk; within the Myc^+^ population, the cutoff for pErk positivity is indicated (horizontal line). Arrows depict quantification of % pErk^+^ cells with increasing expression of Csk^Myc^. **(b)** % Erk positivity vs. Csk^Myc^ expression. The apparent IC_50_ for Csk suppression (dotted line) was obtained by fitting to a sigmoidal dose-response curve (Prism software). Due to incomplete suppression of TCR signaling, K222R fits were constrained so the lower baseline and slope mirrored the average fit values from the other samples. **(c)** Representative dose-response curves showing the relative efficacies of transfected Csk^Myc^ constructs (WT, H21I, K43D, or K222R) in suppressing Erk phosphorylation in TCR-stimulated cells. **(d)** Relative IC_50_ values for Csk constructs; Error bars: SEM, n=3 (K222R), 6 (K43D), 21 (H21I), 27 (WT) independently transfected and stimulated samples. Mix-effects analysis was performed in Prism, with Tukey’s test for multiple comparisons ****p<0.0001, ***p=0.0001.

## Discussion

We present the first cellular evidence that Csk can form homodimers that compete with PTPN22 (LYP/Pep) binding. We used structural analysis to design single-amino-acid substitutions in the SH3 domain of Csk that disrupt either homodimer formation (Csk H21I) or PTPN22 binding (K43D) without impairing kinase activity in solution. We also tested the substitution commonly used to block SH3-domain PXXP binding (Csk W47A) and found a secondary effect of impairing Csk homodimer formation. The PTPN22-specific Csk K43D construct will therefore be a useful tool for selective studies of homodimer vs. phosphatase interactions in cell signaling. Together, the H21I and K43D mutations demonstrate that PTPN22 binding and Csk homodimer formation compete for access to the Csk SH3 domain. This may indicate that multivalent recruitment of Csk to adaptors and other binding partners at the plasma membrane controls the residency of phosphatases alongside their membrane-localized substrates.

Future studies of Csk at physiological expression levels and in primary (non-cancer) cells will be necessary for quantitative analysis of homodimer formation. However, even at physiological expression levels Csk is likely at sufficiently high local concentration to drive dimerization^60^. CBP/PAG1 oligomerizes upon phosphorylation^1,4–6^, and the signaling complexes in the early TCR signalosome are highly oligomeric and bridged. In our experiments, moreover, endogenously expressed proteins would recruit and concentrate Csk at the membrane, limiting the effect of Csk overexpression on access to substrate.

The effect of homodimerization on Csk function remains an interesting question. We found no evidence that Csk H21I had impaired kinase activity, but the analysis was performed at a concentration below the likely K_d_ of homodimer formation. Many tyrosine kinases have complex allosteric networks, including Csk, in which the phosphopeptide-bound SH2 domain increases the activity of the kinase^1,5,45^. SH3-SH3 homodimer interactions could similarly influence Csk kinase activity.

Conversely, loss of either homodimerization or PTPN22 binding decreased the ability of overexpressed Csk to suppress Lck/TCR signaling. We do not yet know whether Csk H21I can form homodimers in the highest-expressing transfectants or whether a single molecule of Csk H21I may pair with endogenous Csk (with loss of only one of two histidine residues in the symmetrical interface). Future studies without competition from endogenous Csk will be necessary to define the physiological function of Csk homodimer formation. The role of Csk-bound PTPN22 is debated, and we cannot definitively address this question. However, our observations that loss of homodimerization (with increased PTPN22/Csk interaction) *or* loss of PTPN22 binding impairs Csk function in T cells does not support a Csk-centered gain-of-function model for PTPN22 R620W.

The dual requirement for homodimerization and PTPN22 binding is particularly interesting because the homodimer and PTPN22 are likely incapable of forming a ternary complex. This suggests that different pools of Csk or a kinetic process of swapping one binding partner for the other is required for optimal Csk function. The more severe defect in K222R function and kinase activity relative to Csk H21I and K43D reaffirms that catalytic activity is the dominant requirement for the inhibitory function of Csk. However, homodimer and PTPN22 interactions with Csk may coordinate to maximize suppression of Lck and the TCR pathway.

We speculate that Csk homodimerization could be a key step in cell signaling. At equilibrium and relatively low concentration in the cytosol, the Csk SH3 domain could bind preferentially to phosphatase. In T cells, half of all PTPN22 molecules are associated with Csk, so this would be a substantial proportion of the available phosphatase activity^19^. Upon recruitment to the membrane, the local concentration of Csk would increase due to enrichment in microdomains and CBP/PAG1 oligomerization. Csk homodimers might then outcompete PTPN22 for access to the SH3 domain. Released near its substrates, PTPN22 would then be free to dephosphorylate and block signaling through Lck, ITAMs, and Zap70. In this gain-of-function mode, the catalytically efficient phosphatase could act on many more substrate molecules after dissociation from the longer-lived Csk/Cbp/Lck complex. Alternatively, in a loss-of-function mode, PTPN22 might bind an alternative ligand (for instance, TRAF3, FAK or Pyk2 kinase^6^, a Dok-family adaptor protein7) and lose access to substrates. It is also possible that a secondary effect of the autoimmunity risk allele PTPN22 R620W could be to increase Csk homodimer formation via loss of PTPN22 competition for SH3-domain access. Future studies will be required to address these questions.

Our structural observations also predict a functional difference between PTPN12 and PTPN22 phosphatases, possibly leading to differential regulation of SFK signaling in non-hematopoietic and hematopoietic cells. The PXXP-only PTPN12/Csk interaction, for instance, might be easily outcompeted by homodimer formation at a modest local concentration of Csk. PTPN12 binds to Csk only 1-10x more tightly than would another Csk molecule, and the PTPN12 binding site is completely occluded by the Csk/Csk interface, preventing rebinding. Hem-atopoietic-cell PTPN22, in contrast, might be more difficult to outcompete. PTPN22 binds to Csk 10-100x more tightly than would another molecule of Csk, and part of the PTPN22 binding site remains exposed even when Csk homodimer is formed. This could facilitate PTPN22 rebinding to Csk and require higher local concentrations of Csk for phosphatase release. These dynamics, in turn, could regulate the signaling thresholds or feedback regulation of the SFKs by cell type, by local concentrations of Csk-binding proteins at the plasma membrane, and by Csk expression itself.

In summary, we report that Csk can homodimerize in Jurkat T cells, in competition with the phosphatase PTPN22. We also present the Csk variants H21I and K43D, new tools for uncoupling these binding processes. Csk expression levels, activity, and localization are all important regulators of signaling^42,61–68^ and can be disrupted in autoimmune disease^25^ and cancer^22–24^. The effect of these variables on homodimerization and PTPN22 binding will be interesting avenues for future mechanistic analysis.

## Materials and Methods

### Structural modeling

Structure-guided mutations were designed from models generated in PyMol (Schrödinger, New York, NY). Alignment and modeling of available NMR and crystal structures revealed amino acids that could be substituted to selectively disrupt homodimerization or PTPN22 binding. The homodimer interface was taken from a crystal structure of the isolated SH3 domain of Csk, in which crystal contacts are formed by the dimerization surface (PDB ID: 1CSK)^51^. The PTPN22-Csk interface was modeled as an amalgam of (i) an NMR structure of the SH3 domain from Csk bound to the 3BP1 peptide from PTPN22/Pep (PDB ID: 1JEG)^13^; and (ii) an NMR structure of the SH3 domain of Src kinase bound to the PXXP-containing peptide APP12 (PDB ID: 1QWE)^44^, with a comparably oriented, canonical PXXP-SH3 interaction. Crystal structures of full-length Csk (PDB ID: 1K9A)^52^ and the phosphatase domain of PTPN22/Pep (PDB ID: 2P6X)^54^ were used to verify the placement of the binding surfaces. Figures were generated with Adobe software (San Jose, CA).

### DNA constructs

Full-length cDNA encoding mouse Csk was subcloned into pEF6 vector (Life Technologies) with a C-terminal linker SAGGSAGG^68^ followed by a Myc (EQKLISEEDL) or HA (YPYDVPDYA) epitope tag. The variants H21I, K43D, W47A^46^, and K222R^3^ were created using QuikChange II site-directed mutagenesis (Agilent).

### Cell culture and transfection

The Jurkat T-cell line E6-1 was maintained in RPMI with 10% heat-inactivated fetal calf serum (FCS) with 2 mM glutamine, penicillin, and streptomycin at 0.15 - 1 M cells/ml in vented flasks in a 37°C cell incubator. For transfections, batches of 15 M log-phase cells in antibiotic-free medium were collected, washed, and resuspended in 300 μl medium with 10 μg empty vector DNA and 20 μg total Csk DNA (e.g. 10 μg Csk^Myc^ + 10 μg Csk^HA^). After 15 min incubation at room temperature, cells were transferred to a 0.4 cm cuvet for a 10 ms pulse of 300 V on a BTX Square-Wave electroporator (Bio-Rad). Cells were rested 15 min at room temperature and then transferred to 10 ml antibiotic-free medium and incubated overnight. Viable cells were quantified with a ViCell counter (Beckman)^65,69^. Jurkat cells from our lab (gift from A. Weiss at UCSF) have tested negative for mycoplasma and been verified by STR profiling and TCR expression.

### Immunoprecipitation and blotting

Lysates of transfected Jurkat cells were made by incubating at least 8 M cells 15 min on ice in lysis buffer containing 1% lauryl maltoside, 150 mM NaCl, 0.02% NaN_3_, 0.4 mM ethylenediaminetetraacetic acid (EDTA, Sigma), protease inhibitors (Sigma), and 10 mM Tris, pH 7.5 (Sigma) and then centrifuging at 4°C for another 15 min. Supernatants were mixed 2 h at 4°C with Protein-A-conjugated sepharose beads (GE Healthcare) and anti-Myc antibody (9B11, Cell Signaling Tech #2276). The beads were then washed in lysis buffer and boiled in reducing SDS PAGE sample buffer. Proteins were resolved by SDS PAGE and transferred to an Immobilon P polyvinylidene fluoride or Immobilon-FL membrane (EMD Millipore)64. Membranes were blocked with 3% bovine serum albumin (Sigma), and then incubated with the primary antibodies goat anti-human PTPN22/Lyp (AF3428, R&D Biosciences), anti-Myc Biotinylated antibody (71D10, Cell Signaling Tech #3946), or anti-HA Biotinylated antibody (C29F4, Cell Signaling Tech #5017). Secondary incubations included horseradish-peroxidase-conjugated rabbit anti-goat IgG (H+L) (Southern Biotech 6164-05), Streptavidin-HRP (Cell Signaling Tech #3999), or LI-COR IRDye secondary antibodies (NC9744100, NC9523609, 926-68072) as appropriate, and blots were developed with the addition of Western Lightning enhanced chemiluminescence reagent (PerkinElmer) and imaged on a Kodak Imagestation or LICOR Odyssey. Densitometry was performed using NIH ImageJ software, and statistical analysis was performed in GraphPad Prism. Brightness/contrast corrections were applied uniformly using Adobe Photoshop software; rotation correction was applied after densitometry analysis.

### Protein purification

Full-length mouse Csk (WT, H21I, K43D, and K222R3) were subcloned into a pGEX-6P-3 vector (GE Healthcare) containing an N-terminal glutathione S-transferase (GST) affinity tag. Each construct was transformed into BL21(DE3) cells (Agilent), and expression was induced with 0.2 mM isopropyl-β-D-thio-galactoside (IPTG) at 18°C overnight. The bacterial pellet was resuspended in GST binding buffer, pH 7.4 (phosphate-buffered saline with 5 mM EDTA and 5 mM dithiothreitol (DTT)) and lysed by freeze/thaw, lysozyme treatment, and sonication by a Branson 450 Sonifier. All purification steps were performed on ice or at 4°C, and all columns and proteases were obtained from GE Healthcare. Clarified lysate was passed through a GST Gravitrap column, and, after washing, GST-Csk was eluted at pH 8.0 with 10 mM reduced glutathione, 25 mM Tris, 50 mM NaCl, and 1 mM DTT. The GST tag was cleaved with PreScission Protease (GE Healthcare) overnight. After buffer exchange by concentration and dilution, Csk was further purified by HiTrap Q anion exchange chromatography at pH 8.0 (50 mM Tris, 50 - 1000 mM NaCl, and 1 mM DTT) followed by a Superdex 200 size exclusion column in 100 mM NaCl, 10% Glycerol, and 50 mM Tris, pH 8.0. Purified Csk was flash frozen in liquid nitrogen and stored at - 70°C. Homogeneity and molecular weight of purified proteins were verified by SDS PAGE with colloidal blue staining (Life Technologies) and mass spectrometry. The concentration of purified Csk was determined from its absorbance at 280 nm using a molar absorptivity of 73800 M^-1^ cm^-1^ as calculated by the ExPASy ProtParam tool^55^.

### Size Exclusion Chromatography

Size exclusion chromatography was performed by loading 1 ml (20 mg, 400 μM) purified Csk (WT or H21I) onto a HiLoad 16/60 Superdex 200 column. Retention volumes were estimated from fits to Gaussian curves (Absorbance=amplitude*exp(−0.5*((volume – mean retention volume)/standard deviation)^2)) and the fit mean retention volume was used to calculate partition coefficients from the function (retention volume - void volume) / (column volume - void volume). The elution of blue dextran marked the void volume, and elution of high-salt buffer (measured by conductance) marked the column volume. To relate the partition coefficient to an apparent molecular weight, partition coefficients of standard proteins from the high-molecular-weight gel filtration calibration kit (GE Healthcare #28403842) were plotted against Log(MW) and fit to a linear function using GraphPad Prism software.

### Kinase Activity Assay

The activity of purified Csk was measured using a continuous spectrometric assay: ATP hydrolysis is coupled via pyruvate kinase and lactate dehydrogenase to NADH oxidation, which results in a decrease in absorbance at 340 nm^53,70^. Kinase activity was measured at 30°C with a Molecular Devices Spectramax 340PC spectrophotometer in a 75 μl reaction with final concentrations of 2.5 μM Csk, 55.7 U/ml pyruvate kinase, 78 U/ml lactate dehydrogenase, 0.6 mg/ml NADH, 1 mM phosphoenolpyruvate, 250 μM ATP, 10 mM Tris pH 7.5, 1 mM MgCl_2_, and 1 mM tris(2-carboxyethyl)phosphine (TCEP). The reaction was initiated by adding a final concentration of 250 μM Csk optimal peptide substrate (KKKKEEIYFFF^57^) synthesized by Elim Biopharmaceuticals (Hayward, CA). Negligible background activity was observed if substrate or kinase were omitted from the reaction. Kinase activity was obtained by fitting the initial segment of the decay curve to a linear function. After fitting, the raw data were normalized so that the maximum value was 1 to facilitate visual comparison. Statistical analysis was performed in GraphPad Prism.

### Cell Stimulation and Flow Cytometry

We quantified pErk after C305 treatment as a readout of Jurkat-cell signaling downstream of the TCR and Lck, which is an all-or-none response in each cell^59^. After resting 15 min at 37°C in serum-free RPMI, C305 (Harlan), an antibody against the Vβ8 chain of the T-cell receptor, was added in prewarmed RPMI to a final dilution of 1:5000. After 2 min, signaling was quenched by fixing cells in an equivalent volume of Cytofix buffer (BD Pharmingen). Cells were collected by centrifugation, washed in a fluorescence-activated cell sorting (FACS) buffer of phosphate-buffered saline (PBS) with 2% FCS and 2 mM EDTA. Cells were permeablized with dropwise addition of ice-cold methanol, combined with vortexing and incubation on ice for 30 min. Cells were barcoded with different dilutions of Pacific Blue and Pacific Orange (BD Pharmingen), washed, and combined. To probe pErk and Csk^Myc^ expression, cells were incubated with rabbit anti-pErk (pT202/pY204, 197G2, Cell Signaling #4377) and mouse anti-Myc (9B11, Cell Signaling #2276), followed by allophycocyanin (APC)-conjugated donkey-anti-rabbit IgG (H+L) (Jackson Immunoresearch 711-136-52) and Alexa488-Goat-anti-Mouse IgG (H+L) (Life Technologies A-11029). After washing and resuspension in FACS buffer, data were collected on a BD Fortessa flow cytometer. Compensation was performed using FACSDiva software with unlabeled cells, cells treated with Pacific Blue or Pacific Orange, and cells labeled with anti-human CD45 (APC, 2D1, eBioscience, 17-9459 and A488, H130, BioLegend, 304019). TCR expression on transfected Jurkat cells was assessed by staining with phycoerythrin (PE)-conjugated mouse anti-human CD3 (BD Pharmingen 555333); TCR expression was unaltered by Csk overexpression.

### Apparent IC_50_ of Csk

Flow cytometric data were analyzed with FlowJo software (Tree Star, Inc.). The live cell population was selected by forward/side scatter, and cells expressing Csk^Myc^ were gated using Alexa488 fluorescence. Populations positive and negative for Erk phosphorylation were gated using APC fluorescence, and a histogram was generated with the number of pErk^+^ and pErk^-^ cells within each Csk expression-level bin. Microsoft Excel was used to calculate % pErk^+^ cells in each bin, and the results were plotted using Graphpad Prism software. Data were fit to the Boltzmann sigmoidal function % pErk^+^ = bottom + (top - bottom) / (1 + exp(V50 - [Csk^Myc^]) / slope). The point of inflection (V50) of each fit was used as an apparent IC_50_ for each Csk construct. Csk K222R fitting was complicated by lack of a lower baseline. To estimate IC_50_ values in this case, the bottom value was constrained to 0 and the slope was constrained to −20000, an average value from the unconstrained fits of WT and other Csk transfectants. After fitting, the curves were normalized between 0 and 100% for ease of comparison. The IC_50_ values from each fit are displayed relative to WT, with WT displayed as individual values relative to the average of all values obtained each day.

## Acknowledgements

We thank Haichuan Liu and the UCSF Helen Diller Family Comprehensive Cancer Center Mass Spectrometry Core Facility for technical and analytical expertise. We thank Bianca Lee, Art Weiss, Erik Peterson, Ethan Corcoran, and Nicholas Levinson for valuable discussions.

## Funding

National Institutes of Health grant T32AR007304 (TSF)

National Institutes of Health grant R01AR073966 (TSF)

National Institutes of Health grant T32DA007097 (BFB)

University of Minnesota Foundation Equipment Award E-0918-01 (TSF)

## Author contributions

Conceptualization: TSF

Methodology: TSF

Investigation: BFB, FVS, TSF

Visualization: BFB, TSF

Funding acquisition: BFB, TSF

Project administration: TSF

Supervision: TSF

Writing – original draft: BFB, FVS, TSF

Writing – review & editing: BFB, FVS, TSF

## Additional Information

Competing interests: The authors declare that they have no competing interests.

Data and materials availability: All data are available in the main text or the supplementary materials.

## Notes

### Competing Interest Statement

The authors have declared no competing interest.

### Summary of Updates

updated text and figures

